# The Specific Genetic Markers to Diagnose Xanthomonads Infecting Pepper and Tomato Plants

**DOI:** 10.1101/2025.01.21.634051

**Authors:** Ahmet Can, Abdul-Nasır Abdul-Baasit, Ilayda Nur Belen, Umran Capar, Burak Sener, Ceyda C. Basaran, Serap Melike Sulu, Mustafa Kusek, Ragip Soner Silme, Omur Baysal

## Abstract

We have improved a marker system to examine the genetic diversity in Xanthomonads species by using whole genome sequence analysis and repetitive regions in the bacterial genome. Specifically, we amplified microsatellite regions using PCR, and the resulting fragments were digested with specific enzymes. These fragment profiles served as molecular markers. Consequently, a more precise marker for bacterial identification was developed by combining the microsatellite and RFLP methods to differentiate Xanthomonads species. In Addition, we analysed the ancestral ordering of various *Xanthomonas* species’ genomes available in NCBI using Progressive Mauve Alignment. The data revealed unique collinear regions characteristic of *Xanthomonas* species. These regions were also associated with cut genomic fragments used as markers, enabling the discrimination of *Xanthomonas* species infecting tomato and pepper. We propose that these findings contribute to understanding the genetic diversity of *Xanthomonas* and rapidly diagnosis.

## 1. INTRODUCTION

Bacterial spot (bacterial scab, black spot) disease is caused by several species of *Xanthomonas* in peppers and tomatoes. This disease shows typical symptoms in the form of lesions on leaves and fruits. This situation can lead to fruit breakdown and fruit rot, causing significant crop losses [1-4].

At present, the available chemical control of the disease is restricted to the agents using copper compounds, which have shown also limited effectiveness. Therefore, when an epidemic occurs, it becomes difficult to suppress the disease development. If the disease occurs early in the season, a highly efficient disease control in fields becomes necessary, which is economically costly and often destroys entire farmers’ crop [1]. *Xanthomonas* infects tomato and pepper plants as well as many other plants [5, 6]. Group of former *X. campestris pv. vesicatoria*, the pathogens responsible for bacterial spot disease in peppers and tomatoes, was selected as the model organism for this study.

The bacterial spot disease can be caused by multiple species of the *Xanthomonas* genus. When grown in culture, these species typically appear as yellow, mucoid colonies. In the past, *X. vesicatoria* was believed to be the sole cause of the disease [7], but it was later reclassified as *X. campestris pv. vesicatoria* [8]. According to Jones et al. [9], the *Xanthomonas* species responsible for bacterial spot disease can be categorized into four groups: *X. euvesicatoria* (*X. campestris pv. vesicatoria*), *X. vesicatoria, X. perforans*, and *X. gardneri*. Collectively, these species are known as *Xanthomonads* [1]. At least two other species, *Xanthomonas campestris pv. raphani* [10] and *X. arboricola* were isolated from tomato and pepper plants [11, 12]. Molecular techniques such as PCR (Polymerase Chain Reaction) designed for specific gene regions (e.g. hrp) associated with virulence are no longer sufficient to distinguish pathogens on their own [13]. Although multi-locus sequencing on multidirectional sequences is considered the most reliable detection method, it may still be insufficient to detect genetic variations within a species. In contrast, molecular markers such as microsatellites, RFLP (Restriction Fragment Length Polymorphism) and AFLP (Amplified Fragment Length Polymorphism) have shown promising results in molecular diagnosis of *Xanthomonas* species [14-16]. RFLP has also been used, to some extent, for geographic classification of pathogens [17].

Tandem Repeats (TRs) are short motifs of DNA fragments that repeat in a consecutive manner. These repetitions can be various due to the addition or deletion of motifs. Variable Number Tandem Repeat (VNTR) loci analysis allows us to target multiple loci with high discriminatory power. Originally referred to as mini- and microsatellite DNA (Simple Sequence Repeats), Nakamura et al. [18] introduced a more precise terminology. They designated the variable loci representing a single locus as VNTR loci. Today, both VNTRs and SSRs (Simple Sequence Repeats) serve as crucial molecular targets for genetic analysis. Microsatellites (also known as SSRs, STRs, or SSLPs) are prevalent in prokaryotes and exhibit a high mutation rate, enabling differentiation between different clonal groups of strains. DNA fingerprinting based on SSRs exemplifies the practical application of these mutations. Moreover, these repetitive sequences provide valuable insights into processes such as gene flow, genetic neighbourhood dynamics, and rates of genetic divergence. This study aims to determine the genetic diversity in *Xanthomonads* species isolated from tomato plants and examine intraspecific differentiation by using microsatellite regions obtained from whole genome sequence analysis as putative molecular markers. For this purpose, PCR amplified microsatellite regions, and the resulting fragments were cut with specific enzymes. The profiles of these fragments were then used as molecular markers. As a result, we have improved a diagnosis using specific markers for bacterial identification by combining powers of both microsatellite (SSR) and RFLP methods.

## 2. MATERIALS and METHODS

### 2.1 Survey Studies for the Collection of Samples

Pepper and tomato plants in commercial production areas in the Serik district of Antalya province, Turkey (36°55′N 31°06′E) were assayed for bacterial diseases at 2020-2022. Samples of plants showing signs of bacterial spots were collected and placed in sterile paper bags, labelled and brought to the laboratory in a cold chain for isolation procedures.

### 2.2 Isolation of the Disease-Causing Agent

Plants with typical of the bacterial spot disease lesions were used to cut plant leaves/stems on the border of diseased and healthy tissue. The samples were surface sterilized in 70% alcohol for 5 min and washed with sterile pure water 3 times. Then, they were crushed with pestil in sterile mortars containing 500 μl sterile water. A loopful of suspension was taken from the obtained extract and streaked onto NA (Nutrient Agar; Merck, Darmstadt, Germany) and YDC Agar (Yeast Dextrose Calcium Agar) media. All inoculated Petri dishes were incubated at 25-26°C and the development of bacterial colonies was examined daily. Suspected colonies were streaked on new Petri dishes with YDC agar and examined for purity. Then, the purified isolates were placed in Eppendorf tubes containing 40% glycerol and stored at -20°C to be used within the scope of the study.

### 2.3 Phenotypic Characterization

Bacterial isolates were subjected to standard biochemical and physiological tests, including growth on YDC Agar [19], GYC Agar [20], SX agar [21], and CKTM [22, 23] media and evaluated for colony development and morphology.

Bacterial isolates were also evaluated for levan formation (depending on whether levan forms on sucrose nutrient agar (SNA) medium prepared by adding 5% sucrose to NA medium) [19], oxidase production (Nutrient agar+1% glucose and 1% N;N;N;N’ Tetramethyl-1.4 phenylene diammonium dichloride freshly prepared solution is dripped on the sterile filter paper by taking a full extract from the post-incubation culture of the isolates and according to the resulting color condition) [24], catalase reaction [19], potato pectolytic activity (by infecting potato tubers) [19], tobacco hypersensitive reaction (HR) (on *Nicotiana tabacum* L. Cv. *Samsun* [19], potassium hydroxide test (KOH) [25] and starch hydrolyzation tests [19].

### 2.4 Pathogenicity Test

The pathogenicity of the isolates was evaluated on pepper (*Capsicum annuum* L.) seedlings *in vivo*. In purpose of this pathogenicity test, the pepper seedlings planted in pots containing 1:1 peat:perlite were grown till the stage of 3-4 true leaves, the suspension at a density of 10^7^ cfu/ml prepared from 36-48 hours old pure bacterial cultures was infiltrated under the epidermis from the leaf lower surface. Reference *X. euvesicatoria* strain (GA-2, US) was used as a positive control and sterile pure water was used as a negative control. Inoculated seedlings were placed in the climate chamber and kept under polyethylene bag for 24 hours to ensure high humidity for initial infection process. The symptoms were rated Yes or No. The typical yellow-halo leaf spots formed on the leaves in 7-10 days after inoculation.

### 2.5 Specific Primers Designing and Restriction Maps

For primers development, *X. campestris* pv. *vesicatoria* 85-10 genome (AM039952.1) was downloaded from the NCBI database. Microsatellite regions were selected by analysing repeat rates and sequence lengths in the genome using http://insilico.ehu.es/site. Three microsatellite loci with core motives (TCGCTG)_11_, (CAGGCAGA)_7_, and (CCCCGAGT)_7_ were selected based on presence of HinfI restriction site (G^ANTC) and total size suitable for PCR amplification and RFLP analysis of obtained PRC product (225-251 bp).

The selected microsatellites are listed in the Table 1 and their repeat numbers are indicated according to *X. campestris* pv. *vesicatoria* 85-10 sequence. Primers specific to the selected microsatellites were designed using the Primer3 program (Whitehead Institute for Biomedical Research, US). The designed primers were used for PCR amplification to determine the genetic diversity in *X. campestris* pv. *vesicatoria* and ensure intraspecific differentiation.

**Table 1.**
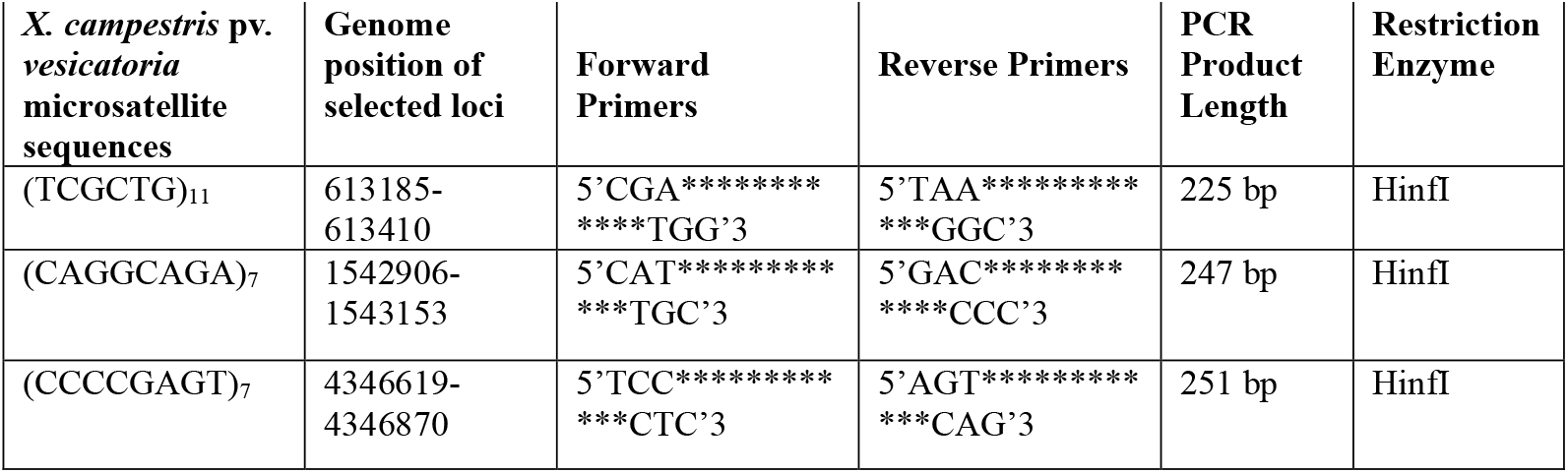
Microsatellite regions, primers and restriction enzymes of *X. campestris* pv. *vesicatoria* strain 85-10.

| *X. campestris* pv. *vesicatoria* microsatellite sequences | Genome position of selected loci | Forward Primers | Reverse Primers | PCR Product Length | Restriction Enzyme |
| --- | --- | --- | --- | --- | --- |
| (TCGCTG)<sub>11</sub> | 613185-613410 | 5'CGA*****<br>****TGG'3 | 5'TAA*****<br>***GGC'3 | 225 bp | HinfI |
| (CAGGCAGA)<sub>7</sub> | 1542906-1543153 | 5'CAT*****<br>***TGC'3 | 5'GAC*****<br>***CCC'3 | 247 bp | HinfI |
| (CCCCGAGT)<sub>7</sub> | 4346619-4346870 | 5'TCC*****<br>***CTC'3 | 5'AGT*****<br>***CAG'3 | 251 bp | HinfI |

#### 2.5.1 Strains Forming PCR Products and Product Sizes

As a result of *in silico* PCR analysis performed with the primers designed for the target microsatellites loci (Table 1) genomes of *Xanthomonas* bacteria that could yield positive results were identified. According to the BLAST results, the sizes of the genomes to which the primer pairs were connected were determined by the *in-silico* PCR test and this information is shown in Table 2, Table 3, Table 4.

**Table 2.** NCBI reference genomes used for primer pair 1.

| Species, strains | PCR | Restriction | Genome position of the loci | GenBank |
| --- | --- | --- | --- | --- |
| *X. campestris* pv. *vesicatoria* 85-10 | 225 bp | 163-55-7 bp | 613185-613410 | CP017190.1 |
| *X. euvesicatoria* pv. *alfalfae* | - | - | - | CP072268.1 |
| *X. vesicatoria* | - | - | - | CP018725.1 |
| *X. perforans* | 195 bp | 133-55-7 bp | 2257558-2257753 | CP116305.1 |
| *X. hortorum* | - | - | - | CP016878.1 |
| *X. euvesicatoria* | 207 bp | 141-59-7 bp | 2299127-2299334 | CP018467.1 |

**Table 3.** NCBI reference genomes used for primer pair 2.

| Species, strains | PCR | Restriction | Genome position of the loci | GenBank |
| --- | --- | --- | --- | --- |
| *X. campestris* pv. *vesicatoria* 85-10 | 247 bp | 159-88 bp | 1542906-1543153 | CP017190.1 |
| *X. euvesicatoria* pv. *alfalfae* | 174 bp | 88-86 bp | 3028172-3028346 | CP072268.1 |
| *X. vesicatoria* | - | - | - | CP018725.1 |
| *X. perforans* | 174 bp | 88-86 bp | 3086070-3086244 | CP116305.1 |
| *X. hortorum* | - | - | - | CP016878.1 |
| *X. euvesicatoria* | 223 bp | 139-84 bp | 1472023-1472246 | CP018467.1 |

**Table 4.** NCBI reference genomes used for primer pair 3.

| Species, strains | PCR | Restriction | Genome position of the loci | GenBank |
| --- | --- | --- | --- | --- |
| *X. campestris* pv. *vesicatoria* 85-10 | 251 bp | 66-33-15-(8)<sub>13</sub>-(7)<sub>4</sub>-5 bp | 4346619-4346870 | CP017190.1 |
| *X. euvesicatoria* pv. *alfalfae* | - |  | - | CP072268.1 |
| *X. vesicatoria* | - |  | - | CP018725.1 |
| *X. perforans* | - |  | - | CP116305.1 |
| *X. hortorum* | - |  | - | CP016878.1 |
| *X. euvesicatoria* | 267 bp | 66-29-15-(8)<sub>14</sub>-(7)<sub>6</sub>-3 bp | 3759280-3759547 | CP018467.1 |

### 2.6 Genomic DNA Isolation

Pure cultures of *Xanthomonas* isolates (a1, a2, i1, i2, u1, u2) were used to determine their colony morphologies. These strains were inoculated into both solid (using Merck nutrient agar) and liquid (using Merck nutrient broth) media. The solid cultures were stored at 25°C for 1 week, while the liquid cultures were incubated for 24 hours. After incubation, genomic DNA samples were isolated using the GeneJET Genomic DNA Purification Kit (Thermo Sci.). A_280_/A_260_ and A_280_/A_230_ ratios and concentrations of DNA samples were measured on a Multiskan™ GO Microplate (Thermo Sci.) spectrophotometer. The obtained genomic DNA samples were run in 1% agarose (Biomax) gel at 90 V constant current for 1 hour and then visualized using the Quantum (Vilber Lourmat) gel imaging system. GeneRuler 100 bp DNA Marker was used as a marker.

### 2.7 Amplification and Cutting of Microsatellite Regions by PCR

In each PCR tube; Ampliqon Taq OptiMix CLEAR 2x Master Mix (Tris-HCl pH 8.5, (NH_4_)_2_SO_4_, 3 mM MgCl_2_, 0.2% Tween, 1.6 mM dNTP mixture, Taq DNA Polymerase) was optimized with 0.4 μM forward and reverse primers (200 nmol HPLC) and 50 ng DNA, reaction components. The final volume was made up to 25 μL with PCR-grade water. Elongation temperatures were calculated as 56°C for the pairs of primers that will replicate the studied regions, and PCR was performed in the form of 30 cycles. After PCR, agarose gel electrophoresis and imaging were performed. The resulting PCR products were cut using the PfeI (FastDigest) enzyme and visualized by agarose gel electrophoresis. The obtained genomic DNA concentration was set at 50 ng/μL. A multiplex PCR was performed using specific primer pairs in the species *X. campestris* pv. *vesicatoria, X. euvesicatoria* pv. *alfalfae, X. vesicatoria, X. perforans, X. hortorum*. These PCR reactions were subjected to the following amplification program: First, an initial denaturation step lasting 5 minutes at 94°C was performed, followed by 30 denaturation cycles lasting 30 seconds at 94°C, an alignment step lasting 64 minutes at 1°C and an extension step lasting 72 minutes at 1°C; finally, a final extension step lasting 7 minutes at 72°C was performed.

The resulting PCR products were analysed by electrophoresis on a 1.5% agarose gel using 1X ethidium bromide intercalating agent for 120 minutes and visualized on Kodak GenLogic 200 photo documentation (NY, USA).

### 2.8 Classification of *Xanthomonads* using molecular markers

Band lengths in the gel images obtained from PCR and cutting products were calculated using the GelAnalyzer 23.1 program. Calculated band lengths were compared with reference genomes from NCBI. Matchs are shown in Table 5.

**Table 5.**
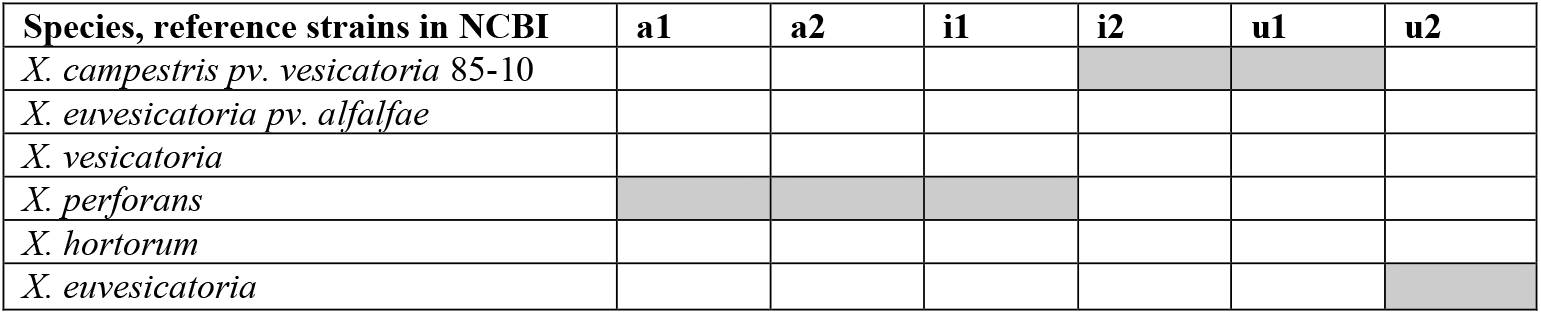
Identification was shown within our strains identified using our primers based on the fragment bands using references strain.

### 2.9 Genomic comparison within *Xanthomonas* subgroups

A total of 6 *Xanthomonas* strain whole genomic sequences available in the NCBI (National Center for Biotechnology Information) database were retrieved (Table 2). These sequences were then analysed using Mauve software from the Darling lab [26]. The objective of this study was to estimate the variation in the gene pool, aiming to identify common genomic regions that represent different subtype diversities based on the origin of the selected whole genome sequences of the strains. Additionally, collinear block analysis was conducted on these 6 genomes to observe a set of loci in different species. These loci, located on the chromosome, was investigated for conservation with the same order or placement in the genome.

## 2. RESULTS

Our analysis shows agarose gel electrophoresis for PCR products amplified using three different primers (primer1, primer2, and primer3) (Figure 1). Under UV light, distinct bands corresponding to specific DNA fragments are visible.

**Figure 1.**
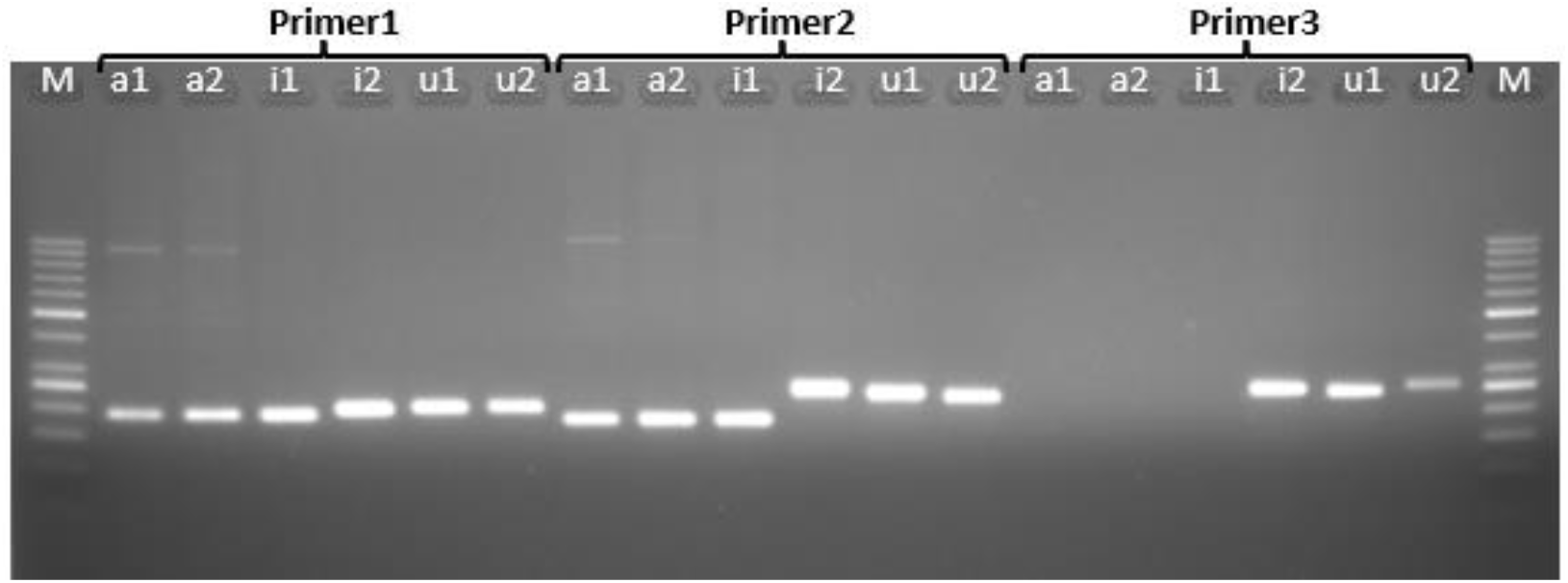
Agarose gel image of PCR products of samples a1, a2, i1, i2, u1 and u2 for primer1, primer2 and primer3

### Primer1 Results

Samples a1, a2, and i1 exhibit a 195 bp band.

Sample i2 shows a 225 bp band.

Sample u1 presents a 225 bp band.

Sample u2 displays a 207 bp band.

### Primer2 Results

Samples a1, a2, and i1 share a 174 bp band.

Sample i2 reveals a 247 bp band.

Sample u1 features a 223 bp band.

Sample u2 maintains a 223 bp band.

### Primer3 results

No bands are observed for a1, a2, or i1 when primer3 is used.

However, in well 17, sample i2 appears with a 251 bp band.

In well 18, sample u1 exhibits a 267 bp band.

The 1–20 bp DNA ladder serves as a reference for fragment sizes. In wells 14, 15, and 16 where Primer 3 was used, there is no band image for bands A1, A2, and i1. However, a 251 bp long band for I2 is found in the 17. well, while a 267 bp long band belonging to the 19. well, is visible for u1 in the 18. well.

We explored the gel electrophoresis results for PCR products (samples a1, a2, i1, i2, u1, and u2) amplified with Primer1, Primer2, and Primer3 (Figure 2). Additionally, we introduced the cutting products obtained using the PfeI enzyme. Here’s the spotlight on those bands obtained after restriction:

**Figure 2.**
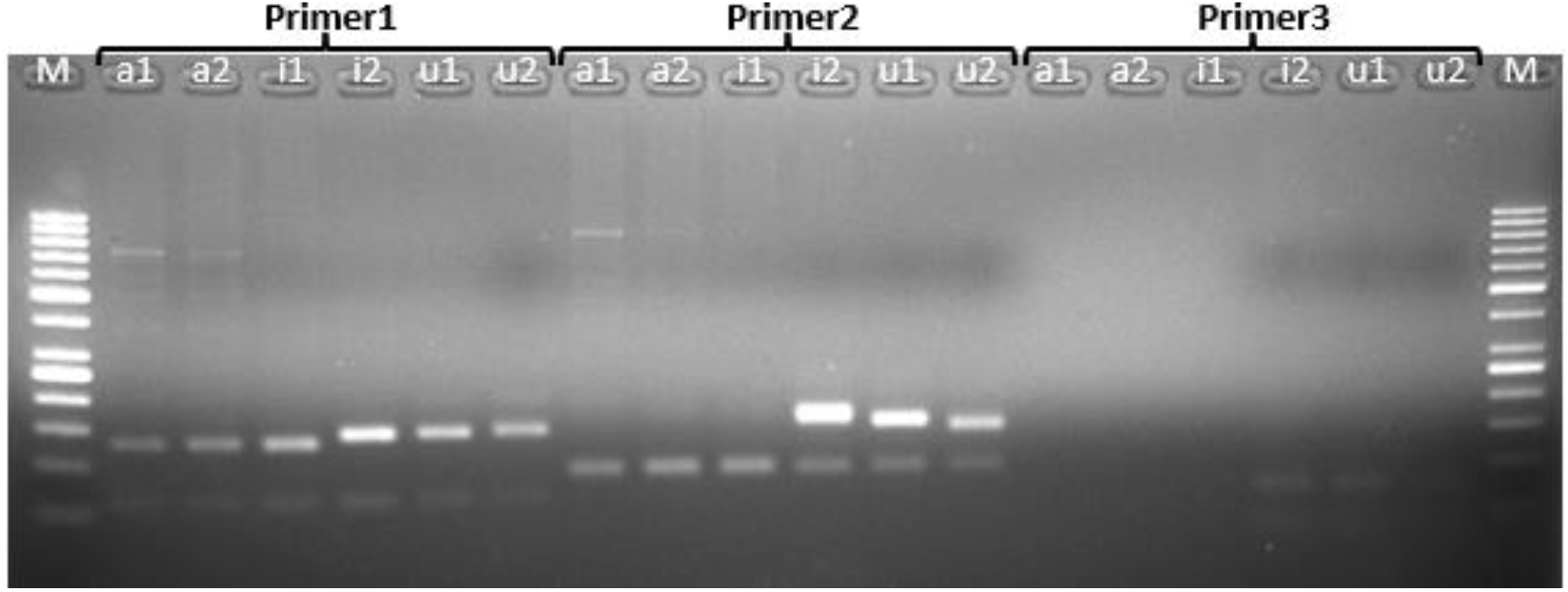
Agarose gel image of PCR products cut with PfeI enzyme.

### Primer1 Cutting Products

Wells 2, 3, and 4 showcase bands at 133 and 55 base pairs.

Wells 5 and 6 reveal bands at 163 and 55 base pairs.

The seventh well flaunts bands at 141 and 59 base pairs.

### Primer2 Cutting Products

Wells 8, 9, and 10 feature bands at 88 and 86 base pairs.

Wells 11 and 12 strut their stuff with bands at 159 and 88 base pairs.

Meanwhile, well 13 boasts bands at 139 and 84 base pairs.

### Primer3 Cutting Products

Wells 14, 15, and 16 remain band-free.

Well 17 grooves to bands at 66 and 33 bp. And in well 18, we’ve bands at 66 and 29 bp.

Depicted data shows the matching of the strains we have with the reference genomes obtained from NCBI (Table 5). According to the band lengths given by the PCR and cutting products of our strains, a1, a2 and i1 showed matched with *X. perforans* strain and i2 and u1 matched with *X. campestris pv. vesicatoria* and strain u2 with *X. euvesicatoria* alone.

Additionally, to observe the confusion in distinguishing species belonging to the *Xanthomonas* genus depending on its genomic profile, genomic comparison analysis was made on the total genome sequences obtained from NCBI. Thus, we have determined the genomic topology difference to understand the reason for the separation efficiency of our designed primes within *Xanthomonas* strains. Genomic comparison studies using the Mauve analysis revealed a genomic comparison among 6 different *Xanthomonas* groups, highlighting the presence of common regions in the bacterial genome (Figure 3). Furthermore, the output from the collinear genome comparison analysis displayed various clusters with sub-clusters that exhibited variation based on the origins of the subtypes according to colinear analysis (Figure 4). Similar clustering patterns were observed when using our selected molecular markers to test *Xanthomonas* strains. These findings indicate that *Xanthomonas* strains exhibit dissimilar genetic diversity, as evidenced by both molecular markers and genomic comparison, in relation to their ancestral genomic clustering.

**Figure 3.**
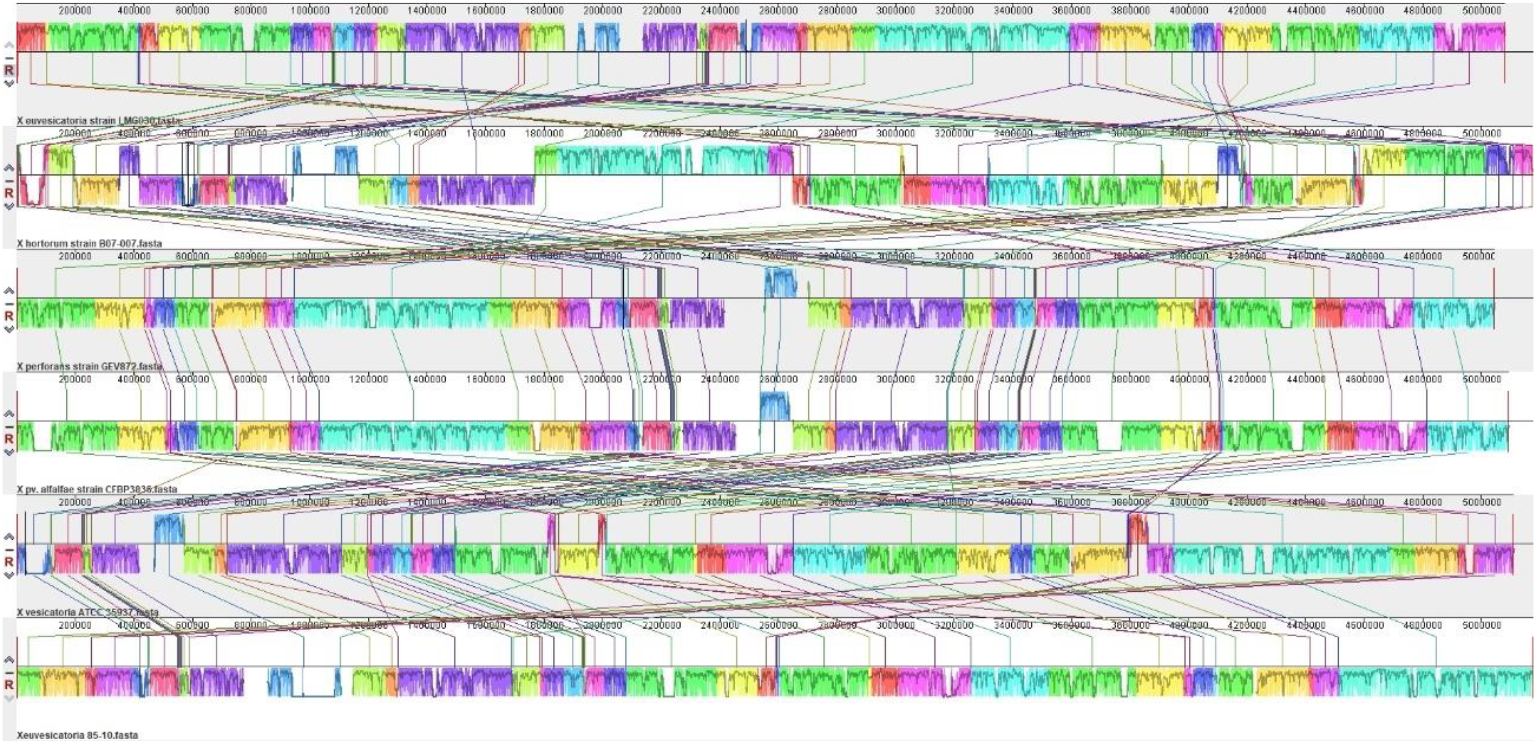
Mauve alignment analysis of 6 different *Xanthomonas* strains genome (CP017190.1, CP072268.1, CP018725.1, CP116305.1, CP016878.1, CP018467.1) accessible in NCBI.

**Figure 4.**
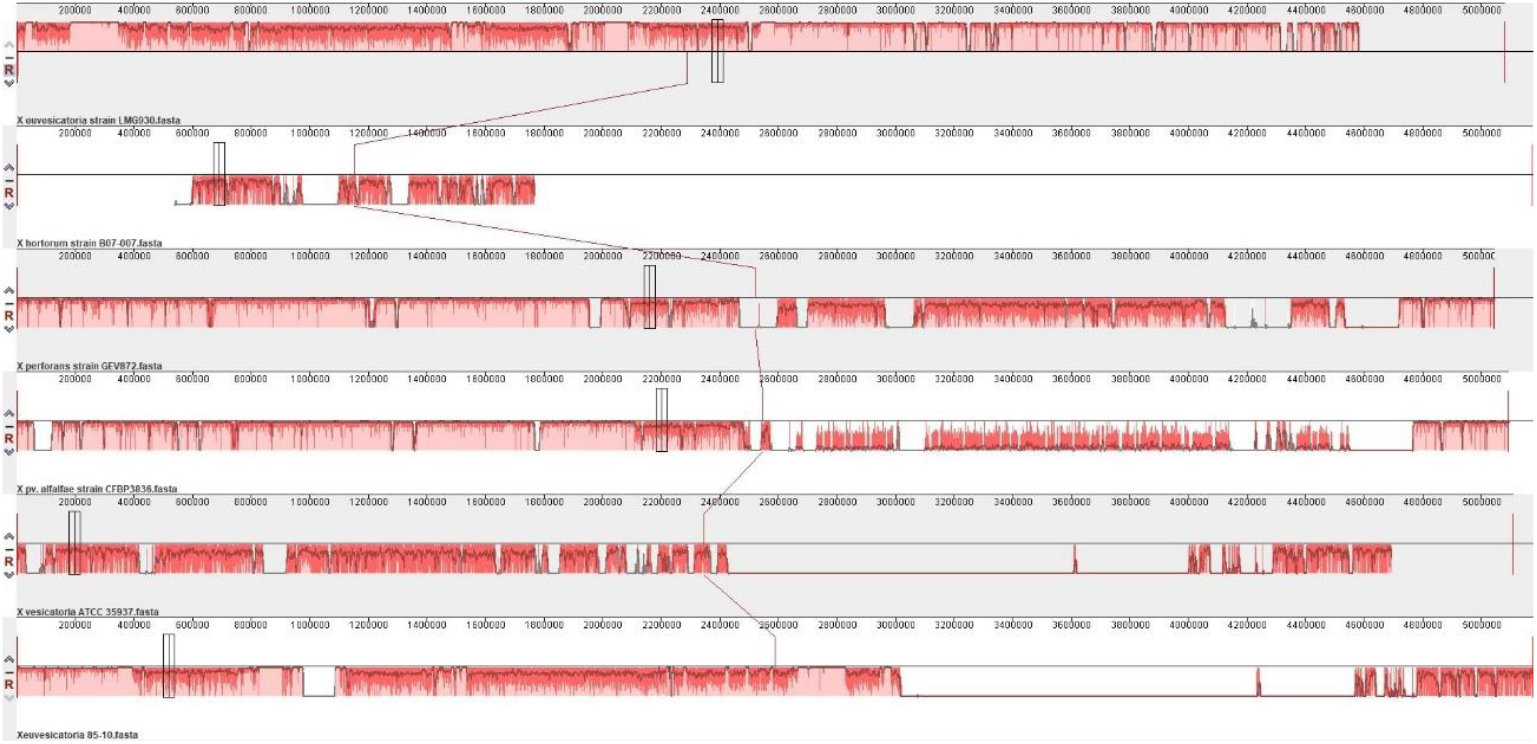
Colinear analysis of 6 different *Xanthomonas* strains genome (CP017190.1, CP072268.1, CP018725.1, CP116305.1, CP016878.1, CP018467.1) available in NCBI indicated various genomics topology.

## 3. DISCUSSION

In this study, a molecular discrimination method developed by combining SSR and RFLP methods was employed to ascertain the genetic diversity of *Xanthomonas* species and to examine intraspecific differentiation. At the molecular level, although methods such as PCR have been employed for pathogen detection, these methods are known to be inadequate for the detection of genetic variation [13]. Consequently, advanced techniques such as versatile sequencing and molecular markers have been introduced. Molecular markers, including RFLP and SSR, have demonstrated considerable potential for differentiating between *Xanthomonas* species [14, 15]. VNTR regions are of great significance in genetic analyses due to their high degree of discrimination [27].

The three microsatellite regions (TCGCTG)_11_, (CAGGCAGA)_7_ and (CCCCGAGT)_7_ played a pivotal role in elucidating the genetic differentiation among *Xanthomonas* species. The microsatellite regions were selected through an analysis with the genome of *X. campestris pv. vesicatoria*. Following a PCR amplification of these regions, the fragments obtained by cutting with specific enzymes were employed as molecular markers. This combination yielded a greater degree of discrimination than that achieved by the methodologies proposed in previous study [27].

The band profiles obtained from the PCR analyses of the isolates were found to be consistent with the band sizes that had been predicted through *in silico* analyses. For example, the amplification of isolates a1, a2 and i1 with primer 1 yielded 195 bp-long bands. A comparison with reference genomes revealed that these bands were consistent with those of the *X. perforans* species. Similarly, isolates i2 and u1 yielded 225 bp long bands upon amplification with primer 1, and these isolates were mapped to *X. campestris pv. vesicatoria*. The PCR results obtained with primers 2 and 3 also corroborated these findings and contributed to the genetic diversity of the isolates and the differentiation between the species.

RFLP analyses permitted a more comprehensive assessment of the PCR products obtained. The band profiles obtained following digestion with the PfeI enzyme served to confirm the successful discrimination of the isolates at the species level. For instance, 133 and 55 bp long bands were observed in isolates a1, a2 and i1, thereby corroborating the genetic similarity of these isolates with *X. perforans*. Conversely, 163 and 55 bp long bands were obtained in isolates i2 and u1, thereby confirming the association of these isolates with *X. campestris pv. vesicatoria*. Isolate u2 gave bands of 66 and 33 bp in length and matched with *X. euvesicatoria*.

In previous studies Klein-Gordon et al. [28] have identified six clusters of *X. perforans* from core gene SNPs that corresponded with phylogenetic lineages. But these SNPs remained limited to only one subgroup. Catara et al. [29] has also suggested the accumulation of genomic information, which give rise to identification of new DNA markers. They will be reason for finding of more sensitive and specific molecular-based methods for the diagnosis and detection of *Xanthomonas* group and diversity. However, Pena et al. [30] have found T3SS and T3SEs are not the only factors that define these lifestyles of *Xanthomonas* groups and SNPs are not sufficient to discriminate of *Xanthomonas* groups and for diversity studies.

In conclusion, the significance of the microsatellite region selection and primer design process was demonstrated in our study. The specificity of the primers to the target regions and the selection of the correct restriction enzymes enabled the clear determination of the genetic differences within the *Xanthomonas* isolates. The combination of SSR and RFLP methods provided high resolution in the discrimination of *Xanthomonas* species, while also contributing to the demonstration of intraspecific genetic diversity. The dissimilarity observed in colinear analysis has supported our finding and indicated why our markers have given rise to segregation in *Xanthomonas* strains. It can be suggested that dissimilarity of colinear regions of various genome could be associated with efficiency of our suggested markers showing amplification in genomes depending on primer sequences considering repetitive regions in genomes. This case could be correlated with host preference differentiation, but we need further detailed studies. These molecular approaches provide a valuable basis for future studies in the monitoring and control of agricultural pathogens and have the potential to make significant contributions to agricultural sustainability. Further studies may investigate the usefulness of this marker system for the discrimination of *Xanthomonas* species in different geographical regions and evaluate its applicability to other plant pathogens. These findings also will contribute to the discrimination of genetic diversity of *Xanthomonas spp* infecting tomato and pepper plants in further studies.

## CREDIT AUTHORSHIP CONTRIBUTION STATEMENT

AC, ABN: Formal analysis; Investigation; Methodology. INB, UC: Methodology. BS: Formal analysis; Investigation; Methodology. CCB: Methodology. SMI: Investigation; Methodology. MK: Formal analysis; Investigation; RSS: Formal analysis; Writing - original draft; Writing - review & editing. ÖB: Conceptualization, Data curation; Formal analysis; Funding acquisition; Investigation; Methodology; Project administration; Resources; Software; Supervision; Validation; Visualization; Writing - original draft; Writing - review & editing.

## ACKNOWLEDGEMENTS

The study has been supported by Muğla Sıtkı Koçman University Scientific Research Projects Coordination Unit (Project Number: 19/095/03/1/2).

## ETHICS APPROVAL STATEMENT

Not applicable

## PATIENT CONSENT STATEMENT

Not applicable

## PERMISSION TO REPRODUCE MATERIAL FROM OTHER SOURCES

Not applicable

## CLINICAL TRIAL REGISTRATION

Not applicable

